# De novo protein design enables targeting of intractable oncogenic interfaces

**DOI:** 10.1101/2025.10.22.683953

**Authors:** Varshika Ram Prakash, Yusuf Najy, Kalel Garrett, Brian F.P. Edwards, Benjamin L. Kidder

## Abstract

Protein-protein interactions (PPIs) involving oncogenic drivers remain among the most intractable targets in cancer biology due to their dynamic conformations and limited accessibility to conventional small molecules. Although antibodies and indirect inhibitors have achieved clinical success against targets such as PD-1/PD-L1 and MYC, challenges persist related to tissue penetration, intracellular delivery, resistance, and incomplete blockade of key interface hotspots. Here, we present DesignForge, an integrated *de novo* protein design framework that combines deep-learning-based structure generation, sequence optimization, and energetic hotspot mapping to create compact miniprotein binders for PPIs. Using this approach, we engineered PD-1 mimetics predicted to disrupt the PD-1/PD-L1 immune checkpoint, designed scaffolds targeting the MYC/MAX dimerization interface, and generated KRAS binders in a manner predicted to occlude RAF interaction. The top designs showed high structural confidence by AlphaFold2, favorable stability metrics, and consistent hotspot engagement identified through MOE-based analyses. Collectively, these results establish DesignForge as a generalizable *in silico* platform for rational design of therapeutic protein binders that extend beyond antibody and small-molecule modalities to systematically target intractable oncogenic PPIs.

## INTRODUCTION

Protein-protein interactions (PPIs) orchestrate essential signaling, regulatory, and structural processes in cancer, yet many oncogenic PPIs remain refractory to conventional small molecules^1^. Unlike enzymes or receptors that contain deep, well-defined catalytic or allosteric pockets, most interaction surfaces are broad, shallow, and conformationally dynamic, optimized for extensive contact areas rather than high-affinity small-molecule binding^1^. Even in cases where pockets exist, such as the nucleotide-binding site of KRAS, they are typically highly conserved, transient, or difficult to access pharmacologically under physiological conditions^2–4^. Antibodies and biologics have extended therapeutic reach to extracellular targets; however, their large size, delivery constraints, and limited intracellular access restrict their applicability to many cancer-associated PPIs^5, 6^. Consequently, new molecular modalities capable of engaging protein interfaces beyond the reach of antibodies and small molecules are highly desirable.

Among known PPI targets, the PD-1/PD-L1 immune checkpoint exemplifies an extracellular interface that has been successfully modulated therapeutically. Antibodies such as nivolumab, pembrolizumab, and others have demonstrated durable antitumor activity across cancer types, making immune checkpoint blockade a central component of oncology therapy^7–9^. Nevertheless, limitations remain, including off-target immune effects, large systemic exposure, cost, and challenges of tissue penetration. Resistance mechanisms, heterogeneity of response, and the need for combination regimens further highlight the unmet need^10, 11^. In particular, antibodies target relatively large surfaces but may not optimally engage defined hotspot residues within the interface. Structural investigations of PD-1/PD-L1 and antibody complexes have revealed diverse blocking mechanisms, including steric overlap with binding surfaces and epitope tuning^12^.

In contrast to PD-L1, MYC represents a nuclear PPI target that has eluded durable pharmacologic inhibition. Aberrant MYC signaling supports proliferation, metabolism alteration, immune evasion, and genome instability^13, 14^. Yet direct targeting of MYC has long been considered “undruggable,” largely because MYC binds DNA and forms heterodimers (e.g. with MAX) via a structurally flexible, shallow interface lacking a clear small-molecule binding pocket^15, 16^. Several small-molecule inhibitors, such as 10058-F4, can disrupt MYC/MAX dimerization, though they generally suffer from limited potency and pharmacokinetic stability^17^. The MYC miniprotein inhibitor Omomyc (OMO-103) has recently entered the clinic, demonstrating safety and early signs of clinical activity in solid tumors^18^. Despite these advances, the MYC/MAX interface remains a challenging and incompletely resolved therapeutic target, motivating alternative *de novo* design strategies capable of achieving precise, high-affinity engagement of this dynamic binding surface^15, 16^.

KRAS represents a cytoplasmic signaling node that, like MYC, was long viewed as intractable but has recently seen progress through allele-specific inhibitors. The advent of covalent small-molecule inhibitors targeting the KRAS G12C mutant (e.g. sotorasib, adagrasib) has shifted that paradigm, demonstrating that select mutant alleles can be drugged^19–21^. However, those inhibitors face significant resistance (both primary and acquired), limited allelic coverage, and challenges beyond G12C^20, 22, 23^. Strategies to stabilize inactive KRAS conformations, disrupt switch regions, or target other mutant alleles remain major gaps.

Recent advances in computational protein design now offer a path to bridging these gaps. Breakthroughs in structure prediction, most notably AlphaFold2, have enabled near-experimental accuracy in modeling protein folds directly from sequence^24^. In parallel, generative modeling approaches, including diffusion-based backbone generation and sequence-structure co-design, are transforming *de novo* binder discovery^25, 26^. Methods that flexibly sample backbone geometries, optimize sequences, and condition on interface constraints have expanded exploration of protein-interaction space, allowing design of scaffolds specifically adapted to defined binding surfaces^27^. When integrated with energetic hotspot mapping, contact-map analysis, and structure-confidence evaluation, such pipelines can systematically yield compact, high-affinity candidates against historically intractable PPIs^28^.

In this work, we integrate BindCraft^29^, a state-of-the-art generative binder design engine, with hotspot identification via MOE’s Protein Contacts module and structural validation via AlphaFold2. This *in silico* framework was applied to three representative oncogenic systems: the PD-1/PD-L1 immune checkpoint, the MYC/MAX transcriptional complex, and mutant KRAS. Across targets, the resulting designs exhibited high predicted structural confidence, thermodynamic stability, and consistent engagement of interfacial hotspots confirmed by contact-map analysis. Collectively, these results demonstrate a generalizable computational pipeline that complements antibody- and small-molecule-based strategies, expanding the therapeutic landscape for previously intractable PPIs in immuno-oncology and oncogene-driven cancers.

## RESULTS

### Hotspot mapping and structural characterization of oncogenic protein-protein interfaces

To delineate the structural and energetic features governing oncogenic protein-protein interactions, we first analyzed representative complexes of MYC-MAX (PDB 1NKP), PD-1/PD-L1 (PDB 5IUS), and KRAS/RAF (PDB 6XHB) using the Molecular Operating Environment (MOE) *Protein Contacts* module (**Fig. 1A-B; Table S1**). This approach quantifies all inter-residue contacts across each interface, assigning both geometric distance and energetic contribution to every residue pair. The resulting profiles highlight residue clusters that make dominant contributions to binding stability, interaction hotspots that define the structural framework for subsequent binder design.

**Figure 1.**
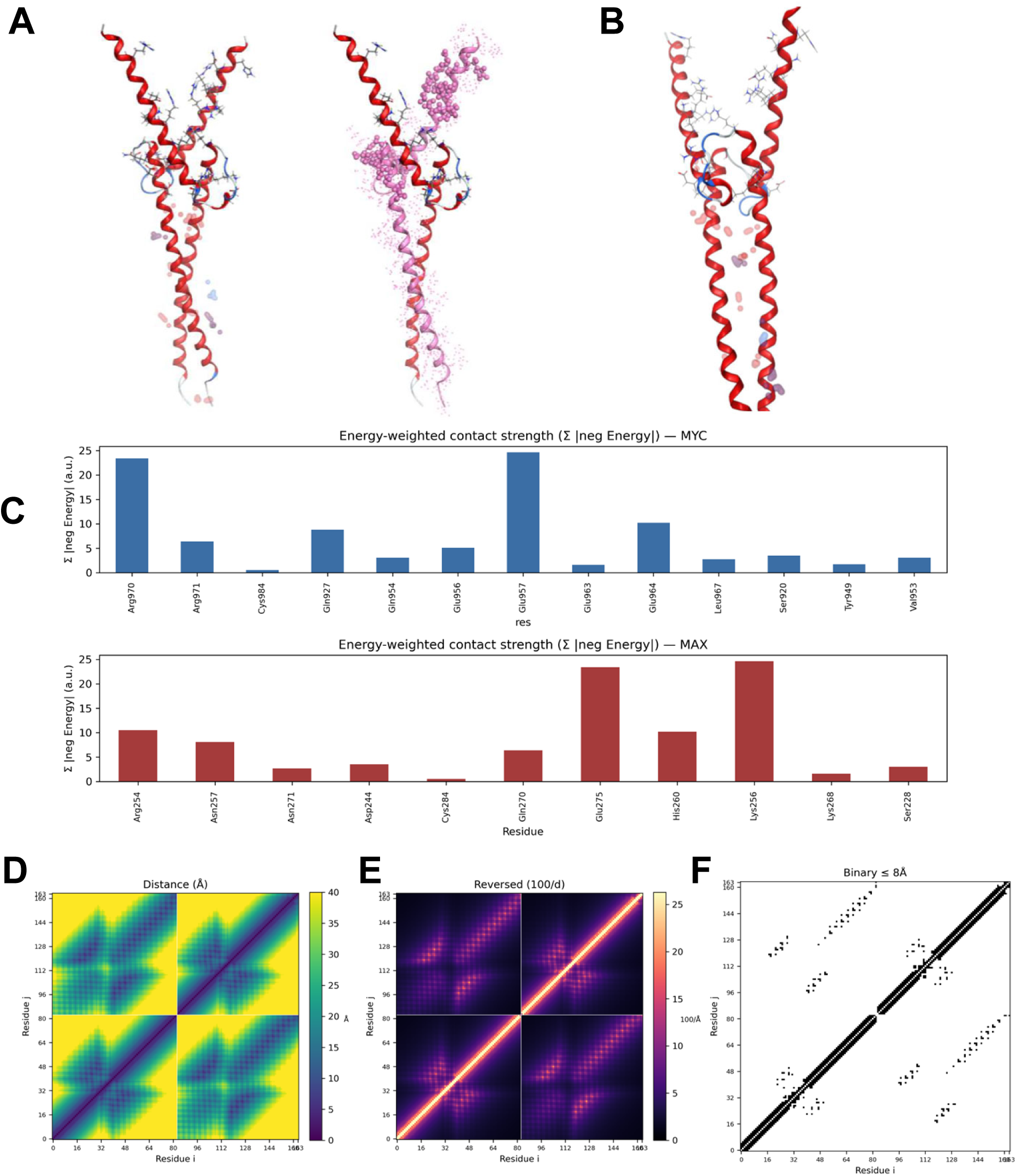
Hotspot mapping and contact analysis of oncogenic interfaces. **(A)** Crystal structure of the MYC-MAX complex (PDB 1NKP) with full heterodimer (left) and isolated MYC chain (right). **(B)** Energetic hotspots on MYC identified by MOE (Molecular Operating Environment) energy decomposition analysis. **(C)** Energy-weighted residue histograms for MYC (top) and MAX (bottom), where bar height represents the sum of absolute negative interaction energies (Σ|Eneg|, arbitrary units). **(D–F)** Contact maps derived from Cα coordinates of the MYC-MAX complex: (**D**) distance matrix (Å), (**E**) reciprocal-distance map (100/d), and (**F**) binary contact map showing residue pairs ≤ 8 Å. Compact interfacial clusters delineate residues mediating the MYC-MAX interaction. Analogous analyses for PD-1/PD-L1 and KRAS/RAF complexes are shown in **Supplementary Figures S1-S2**.

In the MYC-MAX heterodimer, hotspot mapping (**Fig. 1A–B**) highlighted a narrow strip of energetically dominant residues along the coiled-coil interface of the basic helix-loop-helix leucine-zipper region. These residues form a repeating pattern of hydrophobic and electrostatic contacts consistent with prior biochemical studies of MYC/MAX complex formation^30^. The three-dimensional view of the full heterodimer and isolated MYC chain (**Fig. 1A**) emphasizes the exposed helical surface that constitutes the principal binding interface. The MOE energy map (**Fig. 1B**) revealed contiguous patches of favorable contacts primarily involving Gln927, Glu957, Glu964, Leu967, and Arg970 on MYC paired with Arg254, Lys256, His260, Asn271, and Glu275 on MAX (**Table S1**). These hotspots cluster into two continuous bands along the MYC helix, indicating that most of the interaction energy is concentrated within a small subset of residues.

Energy-weighted contact histograms derived from the MOE output (**Fig. 1C**) confirmed that the most stabilizing contributions arise from a small cluster of residues, namely Glu957, Glu964, and Arg970 on MYC engaging Lys256, His260, and Glu275 on MAX. These strong electrostatic contacts account for most of the cumulative negative interaction energy, producing a steep, non-uniform distribution that indicates binding energy is concentrated within a few localized helical turns rather than spread evenly across the interface.

Analogous analyses of PD-1/PD-L1 and KRAS/RAF complexes revealed similarly compact energetic cores (**Supplementary Fig. S1**). In PD-1/PD-L1, the strongest interactions localized around the central FG and BC loops of PD-1, overlapping regions that correspond to known antibody epitopes such as those targeted by nivolumab and pembrolizumab. In KRAS/RAF, contact mapping revealed a dominant high-density cluster centered on the switch I region and adjacent β2-β3 loop of KRAS, which together form the canonical effector-binding surface for RAF. Although switch II undergoes conformational changes during nucleotide exchange, it does not directly engage the RAF interface in this complex. These results reinforce that even in large, seemingly diffuse protein interfaces, binding stability is governed by a restricted set of geometrically and energetically privileged residues.

To complement the energetic analyses, we computed Cα-Cα distance matrices, reciprocal-distance maps, and binary contact matrices using a custom chain-aware Python script (**Fig. 1D-F**). The distance map (**Fig. 1D**) revealed continuous, symmetric interaction bands corresponding to the principal dimerization interface, displaying the periodic lattice typical of parallel helices. The reciprocal-distance representation (**Fig. 1E**) accentuated short-range contacts, separating the high-density interfacial core from peripheral residues. Finally, the binary contact map (**Fig. 1F**) applied an 8 Å cutoff to delineate discrete contact regions, clearly outlining the contiguous inter-helical band that coincides with the energetic hotspots. Together, these geometric representations complement the MOE energy profiles and confirm the spatial organization of stabilizing contacts within the interface.

Collectively, these analyses establish a quantitative description of the molecular determinants that underlie three classes of therapeutically relevant PPIs, nuclear (MYC/MAX), extracellular (PD-1/PD-L1), and cytoplasmic (KRAS/RAF). Across all systems, hotspot residues form compact, energetically concentrated clusters that coincide with structurally accessible and solvent-exposed regions, making them ideal anchor points for *de novo* binder design. The integration of MOE-derived energy mapping with residue-level histograms and distance-based visualization provides a reproducible and generalizable framework for identifying interface features that can be leveraged in the computational generation of selective protein binders.

### Binder design pipeline and structural validation

To extend the hotspot analyses toward binder generation, we applied an integrated computational design workflow combining molecular interface mapping, generative modeling, and structure-based evaluation. The pipeline (**Fig. 2A**) links four sequential stages: hotspot identification in MOE, de novo binder generation with BindCraft, backbone and sequence optimization using RFDesign, and structural validation with AlphaFold2. Together, these modules form a fully in silico framework for creating compact, high-confidence protein binders against the MYC/MAX interface and related oncogenic targets.

**Figure 2.**
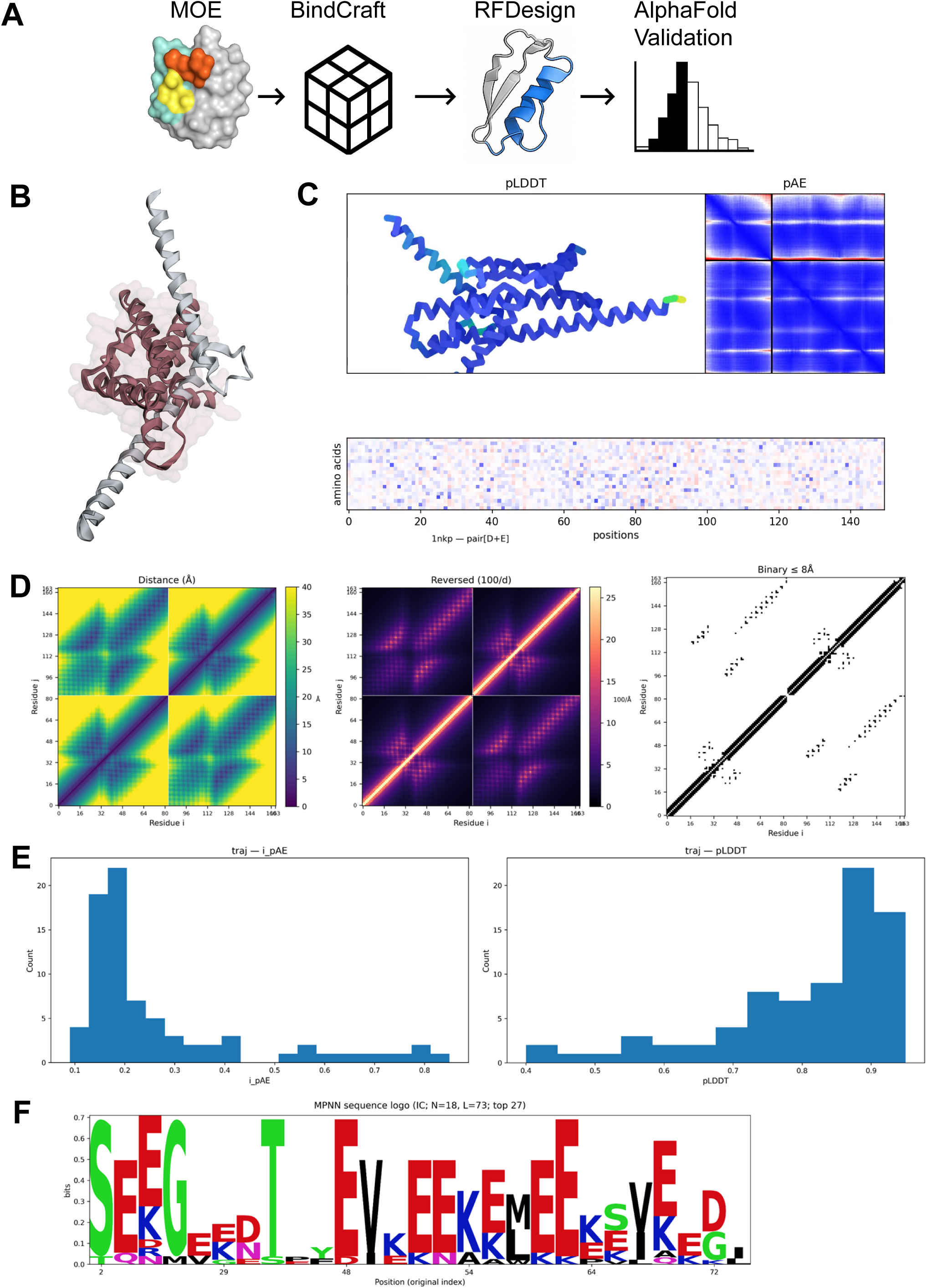
Binder-design pipeline and model validation. **(A)** Schematic overview of the de novo binder design workflow: 1) MOE hotspot analysis, 2) BindCraft scaffold generation, 3) RFDesign refinement, 4) AlphaFold2 structure validation. **(B)** Top-ranked MYC binder model superimposed on the native MYC helix from PDB 1NKP. **(C)** Representative animation frame from AlphaFold2 refinement trajectory showing stable binding orientation with low interface-predicted aligned error (iPAE). **(D)** Distance, reciprocal-distance, and binary (≤ 8 Å) contact maps confirming complementary shape and interface packing between MYC and binder. **(E)** iPAE and predicted local distance difference test (pLDDT) distributions demonstrating high structural confidence and accurate backbone geometry. **(F)** Sequence-logo visualization (information content, bits) highlighting conserved hydrophobic and charged residues within the designed binding interface.

A representative top-ranked binder model generated through this workflow is shown in **Fig. 2B**. The designed miniprotein adopts a compact α-helical fold that aligns along the MYC target helix and mirrors the hotspot geometry identified previously. The interface displays favorable side-chain complementarity near Gln927, Glu957, Glu964, Leu967, and Arg970 on MYC, recapitulating the key energetic contacts observed in the native MYC-MAX complex. The predicted binding orientation positions the designed scaffold to engage both hydrophobic and charged residues across the central coiled-coil region, consistent with the structural requirements for selective MYC recognition.

Trajectory frames from the design refinement process revealed progressive stabilization of the interface and a steady decrease in predicted error. Animation snapshots extracted from the trajectory (**Fig. 2C**) showed that the binder retained a consistent orientation along the MYC helix throughout optimization, with a low interface predicted aligned error (iPAE) value. The representative frame shown exhibits tightly packed helical segments and minimal backbone deviation, indicating a stable, well-defined docking configuration with low structural uncertainty at the interface.

To further assess geometric complementarity, chain-aware distance and contact maps were computed from the final ensemble (**Fig. 2D**). The left panel shows the raw Cα-Cα distance matrix, revealing continuous regions of close proximity between binder and target. The middle panel presents the reciprocal-distance representation (100/d), which enhances visualization of short-range contacts concentrated within the interfacial core. The right panel displays the binary contact map (≤ 8 Å), highlighting a sharply defined band of interacting residues corresponding to the primary MYC binding groove. These complementary visualizations confirm that the designed binder reproduces the contact density and orientation of the native interface while maintaining a compact, self-consistent internal topology.

Beyond geometric validation, we evaluated model confidence and sequence trends among the top-ranked binder designs. Histogram distributions of iPAE and per-residue pLDDT values (**Fig. 2E**) indicated that most trajectories achieved low iPAE (< 0.25) and high pLDDT (> 0.8) scores, signifying high-confidence, well-folded structures. These metrics suggest that the predicted binder-target complexes are geometrically consistent with native-like packing and exhibit minimal uncertainty at the modeled interface.

Sequence variability across the design ensemble was visualized using an MPNN-derived sequence logo (**Fig. 2F**). Strong positional conservation occurred within the central helical region, dominated by hydrophobic (L, F, V) and charged (K, E) residues that mirror the physicochemical pattern of the MYC interface. Peripheral positions displayed greater diversity, reflecting tolerated variability outside the energetic core. The enrichment of leucine and lysine near positions corresponding to Glu957, Glu964, Leu967, and Arg970 supports the residue preferences inferred from the MOE energy mapping and contact-map analyses.

Comparable design-stage analyses for PD-L1 and KRAS binder ensembles are shown in **Supplementary Figure S2**. These models recapitulate the key structural characteristics observed in the MYC designs, including compact α-helical scaffolds aligned along their respective target interfaces (**Fig. S2A**), low predicted interface error and high per-residue confidence (**Fig. S2B**), and dense interfacial contact bands revealed by distance, reciprocal-distance, and binary contact maps (**Fig. S2C**). Histogram distributions of iPAE and pLDDT values (**Fig. S2D**) confirm uniformly high model confidence, while sequence-logo analyses (**Fig. S2E**) highlight strong positional conservation of hydrophobic and charged residues at hotspot-proximal sites. Together, these data demonstrate that the DesignForge workflow yields reproducible, geometrically consistent binders across multiple oncogenic targets.

### Electrostatic surface properties, sequence diversity, and structural confidence of designed binders

Electrostatic potential mapping of the top binder models (**Fig. 3A**) revealed distinct regions of positive and negative surface charge arranged in a pattern complementary to the MYC interface. The interfacial face exhibited a mosaic of alternating acidic and basic patches that align with the charged residues on MYC, suggesting favorable electrostatic complementarity at the binding surface. Across multiple orientations, the designed binder maintained a coherent charge distribution consistent with its intended role as a stabilizing partner to the largely hydrophobic and polar MYC helix.

**Figure 3.**
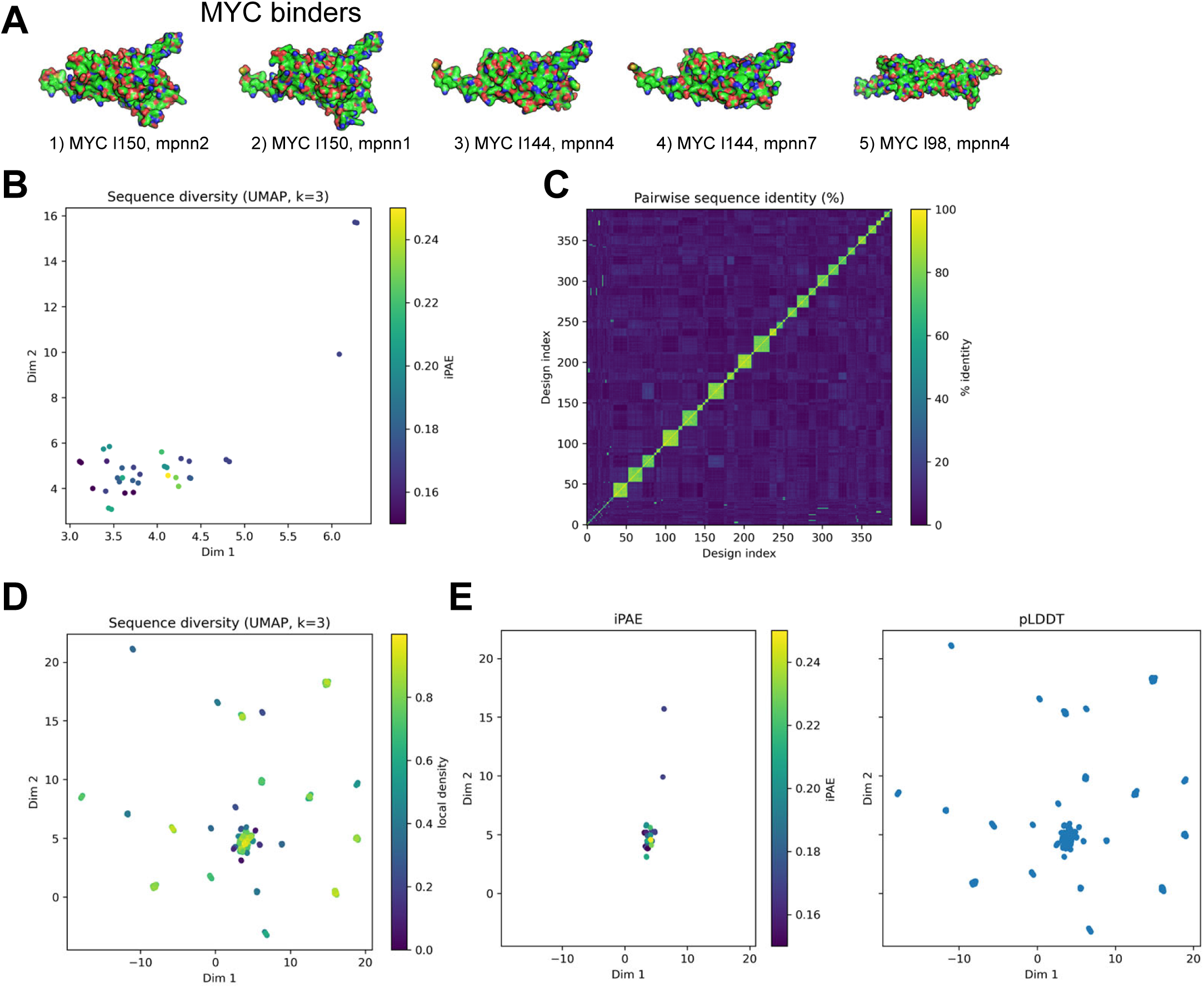
Electrostatics, sequence diversity, and structural metrics of MYC binders. **(A)** Electrostatic surface potential of the top MYC binders (blue = positive, red = negative) showing charge complementarity with the MYC/MAX interface. **(B, D)** UMAP embeddings of binder sequence diversity colored by (**B**) iPAE and (**D**) pairwise sequence identity, generated from the design metrics CSV. **(C)** Pairwise sequence-identity heatmap across top-ranked binders showing strong convergence within the design ensemble. **(E)** Scatterplots comparing iPAE and pLDDT scores for each binder, confirming high-confidence folding and consistent interface predictions. **(F)** ProteinMPNN sequence logo emphasizing recurrent hydrophobic and charged residues at interfacial sites critical for binding.

To assess sequence diversity and structural convergence across the designed ensemble, we projected all variants into a low-dimensional sequence-similarity space using uniform manifold approximation and projection (UMAP). The resulting embedding (**Fig. 3B**) showed that most designs cluster within a compact region, consistent with convergence toward a shared backbone architecture despite underlying sequence variability. A few peripheral points represent more divergent sequences that nevertheless retain low predicted interface error.

Pairwise sequence-identity analysis (**Fig. 3C**) further confirmed substantial sequence diversity, with most designs sharing less than 50% identity. The prominent diagonal reflects unique sequences for nearly all designs, while only small local clusters exhibited moderate similarity. Together, these results indicate that the design ensemble spans a broad and heterogeneous sequence landscape while maintaining a consistent structural fold.

When the analysis was restricted to high-confidence models and colored by iPAE, the UMAP distribution (**Fig. 3D**) revealed a dense central cluster containing the lowest-iPAE binders, surrounded by a broader set of moderately dispersed variants. This pattern indicates that a subset of geometrically consistent, structurally reliable designs emerged from a wider exploration of sequence space.

Relationships between predicted interface accuracy and model confidence were further visualized using iPAE and pLDDT projections (**Fig. 3E**). Most designs clustered at low iPAE and high pLDDT values, underscoring the overall structural reliability of the ensemble. A few outliers exhibited higher error or reduced confidence, while the majority showed stable, well-converged interface geometries.

Collectively, these analyses show that the final MYC binder designs converge toward a shared structural architecture characterized by stable electrostatic complementarity, constrained sequence variability, and high predicted confidence, reflecting both reproducibility and precision of the DesignForge pipeline.

### Structural deviation, interface quality, and sequence diversity among designed binders

To evaluate structural fidelity and sequence convergence of the designed binders, we compared each model to its respective native protein-protein interface and analyzed correlations between backbone geometry, interfacial energetics, and sequence variability. These assessments quantify how accurately the computational designs recapitulate the physical and sequence constraints of their natural counterparts.

Per-residue RMSD analysis (**Fig. 4A**) comparing the top MYC binder to the native MYC interface from the MYC/MAX complex showed close backbone alignment across the helical contact region, with deviations generally within 2-3 Å over most interfacial residues. The global interface RMSD of ∼2.8 Å indicates strong structural fidelity to the target geometry. Local increases near the N- and C-terminal ends correspond to solvent-exposed regions outside the binding interface. The shaded segment denotes the interfacial zone, where the lowest RMSD values coincide with residues contributing the strongest energetic stabilization.

**Figure 4.**
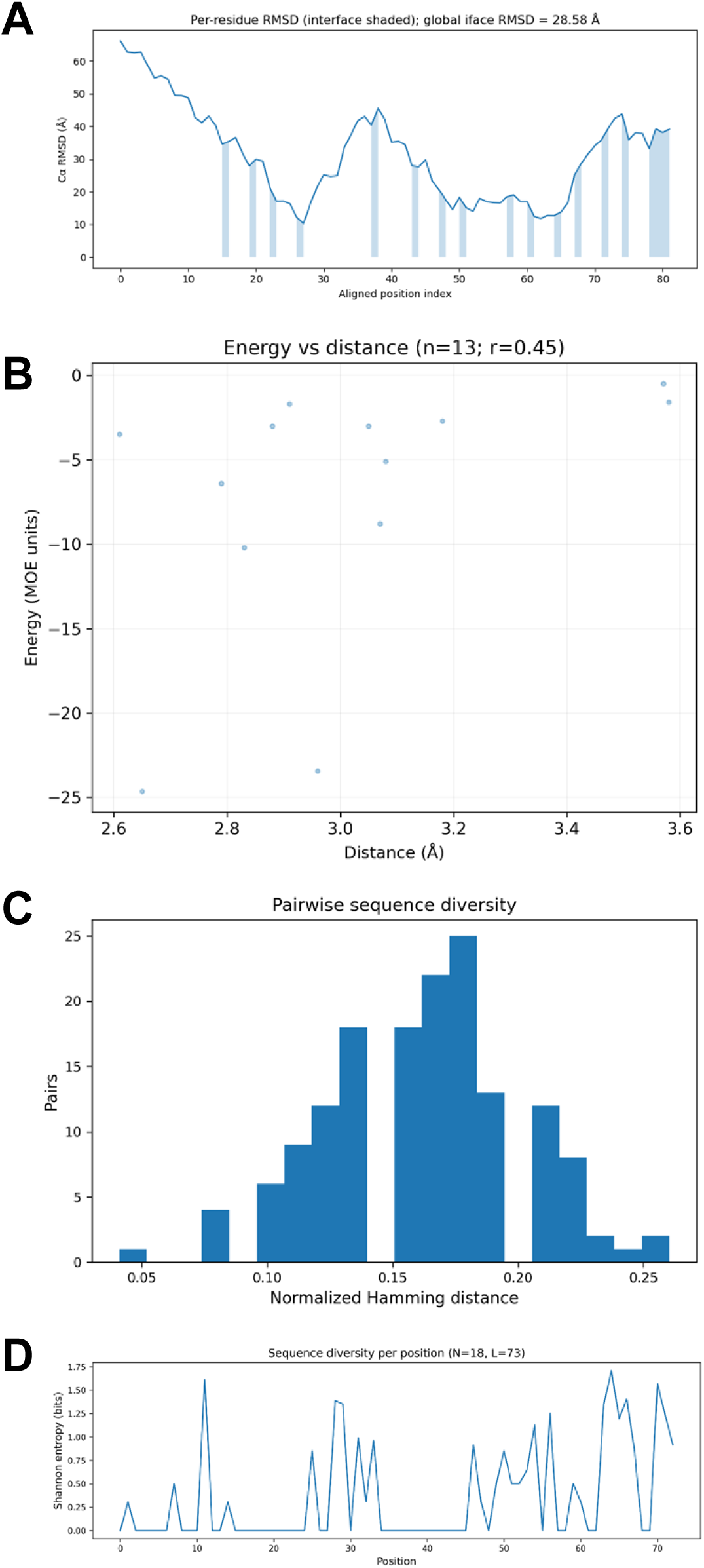
Structural deviation, interface quality, and sequence diversity among designed MYC binders. **(A)** Per-residue backbone RMSD between the top binder mode) and the MYC chain from the MYC/MAX crystal structure (PDB 1NKP). The shaded band indicates the interface region; lower RMSD values within this region denote strong structural agreement with the native motif. **(B)** Scatterplot of interaction energy versus inter-residue distance. **(C)** Histogram of normalized Hamming distances illustrating sequence convergence among designed binders. **(D)** Shannon entropy per residue position, with low-entropy peaks corresponding to conserved interface residues and higher variability at solvent-exposed positions.

The relationship between inter-residue interaction energy and contact distance (**Fig. 4B**) showed a moderate inverse correlation, consistent with residues in closer spatial proximity forming more favorable interactions. This trend reflects physically realistic side-chain packing and effective optimization of non-bonded contacts within the designed interface. Sequence-space analysis demonstrated that the ensemble of designed MYC binders converged toward a narrow set of related variants. The histogram of normalized Hamming distances (**Fig. 4C**) showed a tight distribution consistent with modest sequence diversification around a dominant motif. This limited variability suggests selective optimization toward a stable fold while preserving a small degree of flexibility for local sequence adaptation.

Entropy mapping across aligned sequences (**Fig. 4D**) delineated position-specific variability within the design ensemble. Peaks in Shannon entropy corresponded to loop or solvent-exposed regions, whereas interfacial residues showed minimal entropy, indicating strong conservation driven by structural and energetic constraints. These conserved positions coincide with the MYC hotspot residues identified by MOE contact analysis. These findings reinforce that sequence convergence reflects the physical determinants of binding stability.

To determine whether these design principles generalized across different target systems, we applied the same electrostatic, structural, and sequence-diversity analyses to the PD-1/PD-L1 and KRAS/RAF binder ensembles (**Supplementary Fig. S3-S4**).

For PD-1/PD-L1, Coulombic electrostatic potential maps (**Fig. S3A-B**) revealed contiguous patches of complementary positive and negative charge across the binder-PD-L1 interface, supporting favorable electrostatic complementarity. UMAP embeddings of the design library (**Fig. S3C, E**) showed clustering of low-iPAE, high-confidence binders, consistent with structural convergence toward compact, reproducible folds. Pairwise sequence-identity matrices (**Fig. S3D**) indicated moderate overall sequence similarity, consistent with limited diversity across the PD-1 binder ensemble. When considered together with positional conservation patterns from sequence-logo analyses (**Fig. S2E**), these results suggest that most sequence variability occurs outside the core interface, reflecting focused optimization around the PD-1 epitope.

For PD-L1 and KRAS binders, per-residue RMSD profiles (**Fig. S4A**) revealed broader structural variation relative to their native interface templates, with global interface RMSDs of approximately 22 and 24 Å, respectively. Despite these large overall deviations, both profiles displayed periodic low-RMSD regions within the interface-shaded zones, indicating partial geometric correspondence along the target-contacting helices. The oscillating RMSD patterns suggest that while the designed binders adopt distinct global folds, they maintain localized structural alignment at interfacial residues critical for binding.

The energy-distance relationships (**Fig. S4B**) showed weak or negligible correlations, indicating that favorable interaction energies occur over a broad range of contact distances rather than following a strict inverse trend. Sequence-diversity analyses (**Fig. S4C-D**) revealed substantial variability across the ensembles, with uniformly high positional entropy and broad pairwise Hamming-distance distributions. These results suggest that while the designed binders share a consistent overall topology, they explore a wide range of sequence solutions compatible with the target interface geometry.

The energy-distance relationship (**Fig. S4B**) demonstrated that residues in closer proximity form more favorable interactions. Sequence-diversity metrics (**Fig. S4C-D**) revealed moderate variation across the ensemble with low positional entropy at interface-defining residues, consistent with strong conservation within the switch-region core.

Collectively, these cross-target analyses show that the designed binders for MYC/MAX, PD-1/PD-L1, and KRAS/RAF converge toward geometrically consistent, electrostatically complementary scaffolds characterized by reproducible folding, stable interfacial packing, and high predicted confidence.

## DISCUSSION

This work demonstrates a generalizable, hotspot-guided strategy for de novo mini-protein binder design that spans three mechanistically distinct PPIs: the nuclear MYC/MAX dimer, the extracellular PD-1/PD-L1 checkpoint pair, and the cytoplasmic KRAS/RAF signaling node. Using *DesignForge*, our integrated pipeline combining MOE-based energetic profiling, RFdiffusion backbone generation, ProteinMPNN sequence design, and AlphaFold2 validation produced compact scaffolds that (i) recapitulate near-native backbone geometry at interfacial residues, (ii) exhibit physically coherent energy-distance relationships, and (iii) converge to low-entropy sequence solutions at key hotspot positions. This pattern of sequence convergence likely arises from intrinsic structural constraints, specifically, the limited number of residue combinations compatible with the dense packing and electrostatic complementarity characteristic of high-affinity interfaces, rather than from restricted sampling by the design algorithm.

DesignForge builds directly on recent advances in generative and predictive protein modeling. RFdiffusion generates backbones that satisfy functional constraints while sampling diverse topologies, supporting binder, oligomer, and motif-scaffolding applications within a unified framework^25^. ProteinMPNN then selects sequences predicted to encode those backbones with high fidelity^31^, improving foldability across both native and de novo structures. AlphaFold2 provides structure-confidence metrics (pLDDT, PAE/iPAE) that correlate with experimental success across multiple design studies^24, 32^. The hotspot-driven design strategy extends deep-learning-based functional-site scaffolding approaches, embedding key interfacial residues within compact, energetically optimized folds^33^. Together, these components convert residue-level energetic maps into compact, testable protein scaffolds.

### MYC/MAX

MYC remains a high-value but historically “undruggable” target. The clinical debut of the Omomyc-derived miniprotein (OMO-103) showed first-in-human feasibility for MYC/MAX disruption^18^. Our de novo binders, which recapitulate the MYC/MAX helical-zipper geometry and concentrate conservation at MOE-defined hotspots, align with the emerging view that compact miniproteins can effectively engage transcription-factor PPIs while maintaining favorable biophysical properties. Whereas OMO-103 is MYC-derived, our designs originate from distinct scaffolds that provide an orthogonal starting point for optimizing specificity, stability, and potentially immunogenicity profiles.

### PD-1/PD-L1

Structural and biophysical studies have mapped the PD-1/PD-L1 interface and defined the epitopes targeted by therapeutic PD-1 antibodies such as pembrolizumab and nivolumab^34, 35^. Multiple analyses highlight PD-1 loop regions, particularly FG and BC, as dominant antibody-binding hotspots, consistent with our ensemble results showing low RMSD and high conservation adjacent to these motifs^36–38^. The observed electrostatic complementarity and sequence convergence at loop-proximal positions mirror key features of productive PD-1/PD-L1 blockade interfaces.

### KRAS/RAF

The KRAS-RAF interaction is mediated primarily by the RAF Ras-binding domain (RBD) engaging the switch I region and adjacent β2-β3 loop of KRAS. Structural studies (e.g., PDB 6XHB and related studies^39, 40^) have shown how these extended contact surfaces stabilize signaling assemblies. In our models, per-residue RMSD profiles (**Fig. S4A**) indicated large overall deviations relative to the native complex, yet with localized low-RMSD segments along the switch I-proximal effector-binding surface that preserve its geometric alignment. Energy-distance plots (**Fig. S4B**) revealed weak or negligible correlations, suggesting that favorable contacts occur over a range of spatial separations rather than strictly following an inverse energy-distance trend. Sequence-diversity analyses (**Fig. S4C-D**) showed broad Hamming-distance distributions and moderate-to-high positional entropy, reflecting extensive sequence exploration across the ensemble. Collectively, these results suggest that while the designed KRAS binders adopt distinct global folds, they retain local interface features consistent with engagement of the switch-region surface.

Three quantitative metrics underpin confidence in our designed binders: (i) structural proximity, reflected by low per-residue and interface RMSD values; (ii) physicochemical realism, evidenced by the expected inverse relationship between inter-residue energy and distance, indicating well-packed side chains; and (iii) sequence convergence, characterized by low positional entropy and narrow Hamming-distance distributions that capture reproducible folding constraints. Collectively, these features align with key predictors of success observed in recent experimental validations of RFdiffusion- and ProteinMPNN-based designs^24, 25, 31^.

Two key conclusions emerge from this study. First, hotspot-aware generative design generalizes across structurally and chemically distinct protein-protein interfaces, yielding compact scaffolds whose electrostatics complement the target surface and whose sequence landscapes converge at functionally critical residues. Second, DesignForge introduces quantitative, human-interpretable diagnostics, including RMSD, energy-distance relationships, and sequence-entropy metrics, that enable rational triage of computational candidates before experimental screening and provide explicit parameters for iterative refinement.

In the case of MYC/MAX, where clinical activity has now been demonstrated for a miniprotein class (OMO-103^18^), our results suggest that de novo scaffolds could complement or even surpass derivative constructs through enhanced specificity and stability. For PD-1/PD-L1 and KRAS/RAF, where structural determinants and loop or switch flexibility are well characterized^34, 35, 39^, the observed interface alignment and conservation patterns mirror known contact topologies and motivate targeted experimental validation.

## Conclusion

Integrating hotspot mapping with generative design enables the creation of binder ensembles that converge on sequence-constrained solutions across three representative oncogenic protein-protein interfaces. By anchoring designs at densely interacting energetic cores and validating structural plausibility through AlphaFold-based confidence metrics, this strategy provides a generalizable blueprint for rapid and interpretable in silico binder design.

## MATERIALS & METHODS

### Hotspot mapping and interface analysis

Crystal structures of MYC/MAX (PDB 1NKP), PD-1/PD-L1 (PDB 5IUS), and KRAS/RAF (PDB 6XHB) were analyzed in the *Molecular Operating Environment* (MOE, Chemical Computing Group) to identify energetic hotspots that define key binding determinants. Each complex was imported as a biological assembly, and chains were assigned as receptor and ligand for *Protein Contacts* analysis. The resulting residue-pair tables contained atomic distances and interaction energies, which were exported as CSV files for downstream processing.

Residue-level interaction profiles were computed with custom Python utilities that aggregate the cumulative magnitude of negative interaction energy per residue. Contacts with interaction energy ≤ −1.0 kcal mol⁻¹ and interatomic distance ≤ 3.5 Å were classified as strong. Energy-weighted histograms were plotted to identify dominant residues on both partners that contribute disproportionately to interface stability.

Three complementary geometric representations, Cα-Cα distance, reciprocal-distance (100/d), and binary (≤ 8 Å) contact matrices, were generated to visualize the spatial organization of the interface. These maps confirmed that most favorable contacts in MYC/MAX, PD-1/PD-L1, and KRAS/RAF occur within narrow helical or loop regions, consistent with discrete hotspot clustering.

### Binder design pipeline and structural validation

The design workflow integrated MOE-derived hotspot coordinates with a diffusion-based binder generation framework (BindCraft) and RFDesign for backbone and sequence optimization. Hotspot coordinates served as geometric constraints for conditioning the generative model. For each target, thousands of candidate backbones were sampled, and the 100 highest-confidence designs, meeting predefined thresholds of mean pLDDT ≥ 80 and iPAE ≤ 15 Å² after ProteinMPNN sequence optimization, were retained. Structural confidence and interface accuracy were evaluated using AlphaFold2, and final models were rendered in PyMOL to assess binder orientation and convergence.

Sequence characteristics across the ensemble were summarized using positional frequency logos derived from MPNN output alignments. These logos revealed enrichment of hydrophobic and charged residues at interfacial positions and greater variability at solvent-exposed residues, reflecting selective pressure for electrostatic complementarity and core packing stability.

### Electrostatic, sequence, and structural-metric analyses

Electrostatic surface potentials for top binders were computed using Poisson-Boltzmann calculations within PyMOL and rendered on solvent-accessible surfaces to assess charge complementarity with the target protein.

Ensemble-level metrics were derived from a unified design metrics file containing AlphaFold2 and structural descriptors. Sequence embeddings were projected into a three-dimensional space using Uniform Manifold Approximation and Projection (UMAP; k = 3) to visualize sequence and structure diversity. Pairwise sequence-identity matrices were computed to quantify convergence, and density clustering was used to identify low-iPAE, high-confidence subsets.

Scatterplots of iPAE versus pLDDT confirmed that most binder trajectories clustered tightly at low error and high confidence, indicating consistent model reliability. Pairwise sequence-identity matrices revealed moderate overall similarity indicating partial sequence convergence toward a common structural fold.

### Structural deviation, model quality, and sequence diversity

Backbone deviation of each designed binder relative to its native protein–protein interface was quantified using per-residue root-mean-square deviation (RMSD). For MYC designs, the top model was structurally aligned to the MYC chain from the MYC/MAX complex (PDB 1NKP). RMSD values were computed for backbone atoms (N, Cα, C, O) after least-squares superposition and plotted along the aligned residue index. The shaded region in **Figure 4A** denotes residues comprising the interface as defined by MOE contact analysis (distance ≤ 3.5 Å). Equivalent procedures were applied to PD-1/PD-L1 (PDB 5IUS) and KRAS/RAF (PDB 6XHB) complexes to generate per-residue RMSD profiles shown in **Supplementary Figures S4A**.

### Interface quality and energy-distance correlation

Inter-residue interaction energies and centroid-to-centroid distances were extracted directly from the MOE Protein Contacts tables used for hotspot mapping. For each residue pair, the mean interatomic distance and total interaction energy (kcal mol⁻¹) were plotted as a scatter distribution to assess packing quality. The Pearson correlation coefficient (r) from linear regression quantified the expected inverse relationship between contact distance and interaction energy. Correlation values were observed across designed interfaces, reflecting realistic side-chain packing and physically coherent non-bonded interactions.

### Sequence diversity metrics

All designed binder sequences were aligned using pairwise Needleman-Wunsch global alignment implemented in Python.

Normalized Hamming distances were computed for every sequence pair to measure global sequence divergence. The resulting distribution histograms represent overall sequence diversity for the ensemble (**Fig. 4C**; **Supplementary Fig. S4D**).

Positional sequence variability was quantified using Shannon entropy (Hₛ) calculated from multiple-sequence alignments according to:

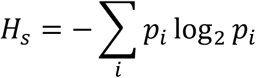

where *pᵢ* is the observed frequency of residue *i* at each alignment column. Entropy values were plotted as a function of residue position (**Fig. 4D**; **Supplementary Fig. S4D**) to identify conserved and variable regions. Interface residues consistently showed minimal entropy, while loop and solvent-exposed regions displayed moderate variability.

### Cross-target comparative analyses

The same RMSD, energy-distance, and sequence-diversity pipelines were applied to the PD-L1 and KRAS binder ensembles, which are presented side-by-side in **Supplementary Figures S3 and S4**. Each figure includes results for both systems to enable direct visual comparison of structural and sequence-based metrics. For PD-L1 binders, alignments were performed to the PD-1 receptor chain using Cα superposition, whereas for KRAS binders, alignments were based on the switch I-proximal effector-binding surface of KRAS. All analyses used identical thresholds and plotting parameters to ensure comparability across target classes.

## DECLARATIONS

### ETHICS APPROVAL AND CONSENT TO PARTICIPATE

Not applicable

### CONSENT FOR PUBLICATION

All authors have read and approved the final version of this manuscript.

#### Statistics and Reproducibility

All analyses were performed entirely in silico using standardized computational workflows to ensure reproducibility. For each target (MYC/MAX, PD-1/PD-L1, and KRAS/RAF), thousands of candidate backbones were generated with BindCraft until 100 high-confidence designs satisfied predefined quality thresholds (mean pLDDT ≥ 80 and interface predicted aligned error ≤ 15 Å²). Hotspot residues were defined in MOE as residue pairs with interaction energy ≤ −1.0 kcal·mol⁻¹ and interatomic distance ≤ 3.5 Å. Structural fidelity was evaluated using Cα-Cα distance matrices, reciprocal-distance and binary (≤ 8 Å) contact maps, and per-residue RMSD relative to native complexes. Energy-distance correlations were derived from MOE contact data and confirmed physically realistic side-chain packing. Sequence diversity was quantified by normalized Hamming distance and Shannon entropy, and electrostatic complementarity was assessed using Poisson– Boltzmann surface potentials rendered in PyMOL. All stochastic processes were executed with fixed random seeds to ensure deterministic reproducibility.

#### Code Availability

All custom scripts used for contact-map generation, binder design, and figure production are openly available at https://github.com/KidderLab/DesignForge

### AUTHORS’ CONTRIBUTIONS

B.L.K. conceptualized the study, designed and performed the computational work, performed data analysis, generated the computational code, and prepared the manuscript. V.R.P., Y.N., and K.G. assisted with generating candidate binders using BindCraft. B.E. provided access to MOE software.

## ACKNOWLEDGEMENTS

We thank Philipp Huettemann for helpful discussions. We also thank the Wayne State University High Performance Computing Grid (https://www.grid.wayne.edu/) for access to the computational resources that made this work possible.

