## Supplemental Information for "De novo protein design enables targeting of intractable oncogenic interfaces"

Running title: De novo designed binders for intractable oncogenic interactions.

[*]Correspondence:

Benjamin L. Kidder

**Supplemental Figures**

**Figure S1. Comparative hotspot and contact-map analyses of PD-1/PD-L1 and KRAS/RAF complexes.**

**(A-B)** Energy-weighted residue histograms derived from MOE interaction tables for (A) PD-1 versus PD-L1 and (B) KRAS versus RAF. Bar height represents the summed magnitude of negative interaction energies (Σ|Eneg|, arbitrary units), highlighting residues contributing most to interface stabilization. **(C)** Overlay of MOE-predicted energetic hotspots on PD-1/PD-L1 and KRAS/RAF crystal structures, showing spatial clustering of dominant interface residues. **(D-F)** PD-1/PD-L1 inter-chain contact maps computed from Cα coordinates: (**D**) distance matrix (Å), (**E**) reciprocal-distance representation (100/d), and (**F**) binary contact map (≤ 8 Å). The continuous contact band corresponds to PD-1 FG and BC loop regions engaging PD-L1. **(G–I)** KRAS/RAF contact maps generated using identical parameters: (**G**) distance, (**H**) reciprocal-distance, and (**I**) binary (≤ 8 Å) representations. Compact contact clusters align with the KRAS switch I/II regions bound by the RAF Ras-binding domain (RBD).

**Figure S2. Structural, contact-map, and sequence analyses of designed PD-L1 and KRAS binders.**

**(A)** Ribbon representations of the top-ranked PD-L1 and KRAS binder designs, each adopting a compact α-helical scaffold positioned along its respective target interface. **(B)** Representative AlphaFold2 structure predictions for PD-L1 and KRAS binders with per-residue interface-predicted aligned error (iPAE) plots, showing uniformly low error and stable backbone geometry across refinement frames. **(C)** Chain-aware contact maps for both binder complexes: *top*, PD-L1 binder; *bottom*, KRAS binder. Each set shows distance (left), reciprocal-distance (100/d, middle), and binary (≤ 8 Å, right) representations. Both designs exhibit dense interfacial contact bands consistent with native binding topologies. **(D)** Histogram distributions of iPAE and predicted local distance difference test (pLDDT) values across AlphaFold2 trajectories: *top*, PD-L1 binder; *bottom*, KRAS binder. Both profiles indicate high structural confidence and minimal predicted interface error. **(E)** Sequence logos derived from top-ranked ProteinMPNN designs: *top*, PD-L1 binder; *bottom*, KRAS binder. Conserved hydrophobic and charged residues dominate buried interfacial positions.

**Figure S3. Electrostatic, structural, and sequence-level analyses for PD-1/PD-L1 binders.**

**(A)** Electrostatic surface potential of the top PD-L1 binder model (*blue = positive; red = negative*), showing complementary charge distribution relative to the PD-1 interface. **(B)** Electrostatic surface potential of the top KRAS binder model, illustrating charge complementarity and interfacial polarity aligned with the RAF-binding surface. **(C)** Pairwise sequence-identity heatmaps for the PD-L1 (left) and KRAS (right) binder ensembles. **(D)** Uniform Manifold Approximation and Projection (UMAP) embeddings of the complete PD-L1 (left) and KRAS (right) binder libraries, where each point represents a designed sequence colored by iPAE. Low-iPAE, high-confidence designs cluster tightly, indicating reproducible backbone geometries and consistent interface solutions across independent trajectories. **(E)** Scatterplots of iPAE versus pLDDT for PD-L1 (left) and

KRAS (right) binders, demonstrating dense clustering in the high-confidence/low-error regime and confirming robust structural reliability of the top-ranked designs.

**Figure S4. Structural deviation, energetic correlation, and sequence-diversity analyses for KRAS/RAF binders.**

**(A)** Per-residue backbone root-mean-square deviation (RMSD) of the top PD-L1 (left) and KRAS (right) binders relative to their native complexes (PD-1/PD-L1, PDB 5IUS; KRAS/RAF, PDB 6XHB). **(B)** Correlation between inter-residue interaction energy and spatial distance across interface pairs for PD-L1 (left) and KRAS (right) binders. **(C)** Histograms of normalized Hamming distances within each binder. **(D)** Per-residue Shannon entropy profiles.

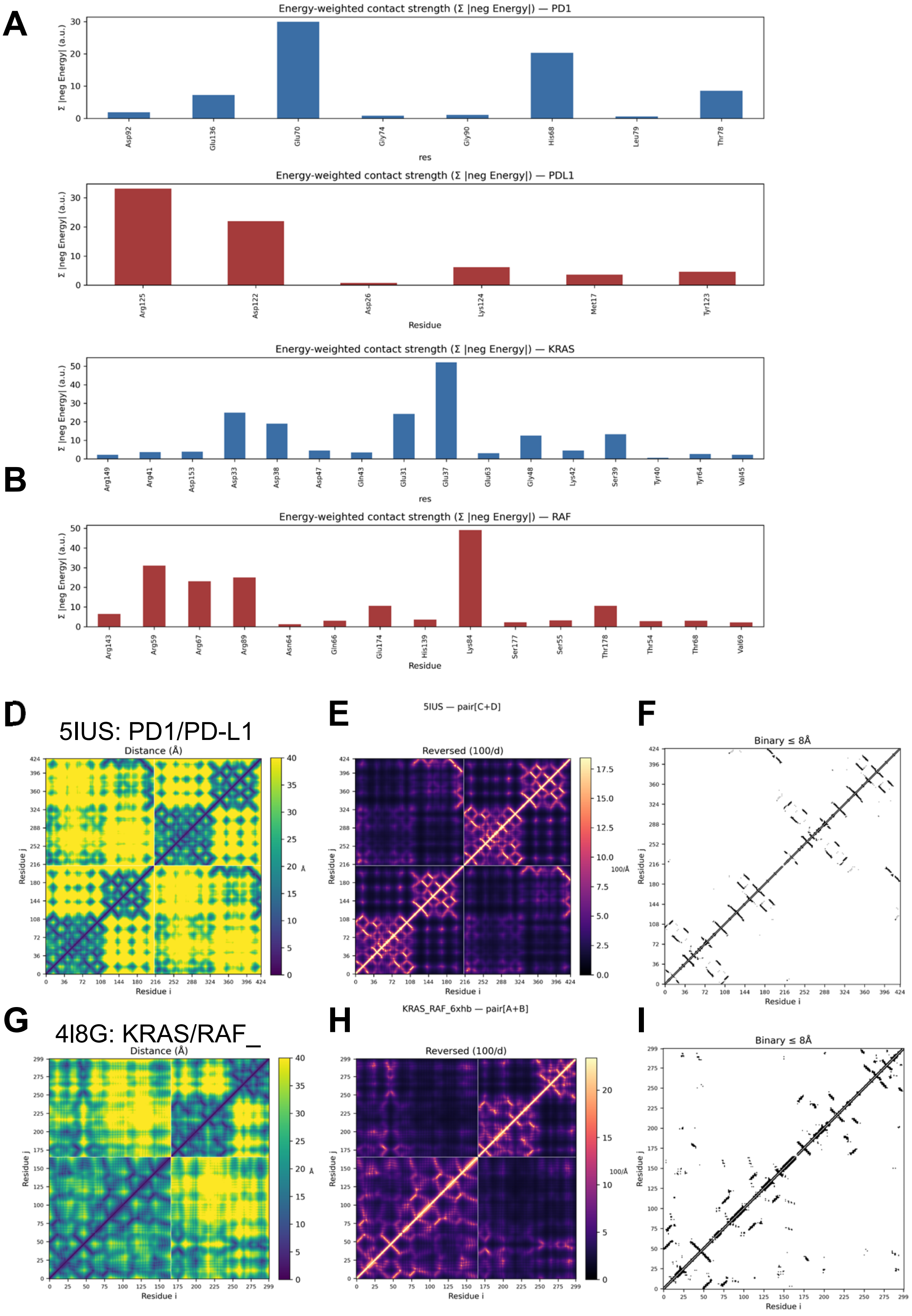
A
Energy-weighted contact strength (Σ |neg Energy|) — PD1
Σ |neg Energy| (a.u.)
Asp92
Glu136
Glu70
Gly74
Gly90
His68
Leu79
Thr78
res
Energy-weighted contact strength (Σ |neg Energy|) — PDL1
Σ |neg Energy| (a.u.)
Arg125
Asp122
Asp26
Lys124
Met17
Tyr123
Residue
Energy-weighted contact strength (Σ |neg Energy|) — KRAS
Σ |neg Energy| (a.u.)
B
Arg149
Arg41
Asp153
Asp33
Asp38
Asp47
Gln43
Glu31
Glu37
Glu63
Gly48
Lys42
Ser39
Tyr40
Tyr64
Val45
res
Energy-weighted contact strength (Σ |neg Energy|) — RAF
Σ |neg Energy| (a.u.)
Arg143
Arg59
Arg67
Arg89
Asn64
Gln66
Glu174
His139
Lys84
Ser177
Ser55
Thr178
Thr54
Thr68
Val69
Residue
D
5IUS: PD1/PD-L1
Distance (Å)
Residue j
Residue i
Å
E
5IUS — pair[C+D]
Reversed (100/d)
100/Å
F
Binary ≤ 8Å
G
4I8G: KRAS/RAF_
Distance (Å)
H
KRAS_RAF_6xhb — pair[A+B]
Reversed (100/d)
I
Binary ≤ 8Å

**Figure S1**

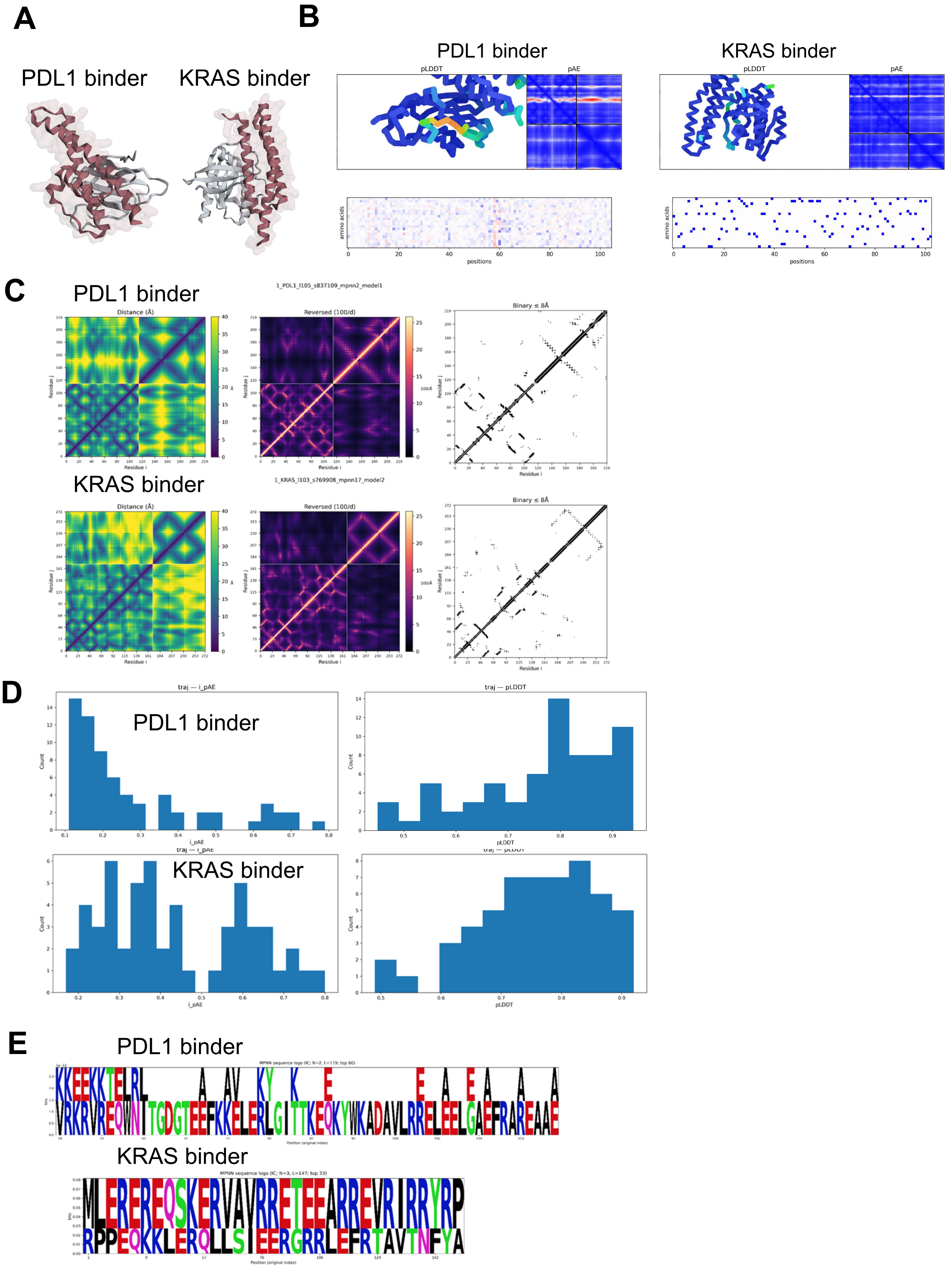
A
PDL1 binder
KRAS binder
B
PDL1 binder
KRAS binder
pLDDT
pAE
amino acids
positions
C
PDL1 binder
1_PDL1_l105_s837109_mpnn2_model1
Distance (Å)
Reversed (100/d)
Binary ≤ 8Å
Residue i
Residue j
KRAS binder
1_KRAS_l103_s769908_mpnn17_model2
D
traj — i_pAE
traj — pLDDT
PDL1 binder
KRAS binder
Count
i_pAE
pLDDT
E
PDL1 binder
KRAS binder
bits
Position (original index)

Figure S2

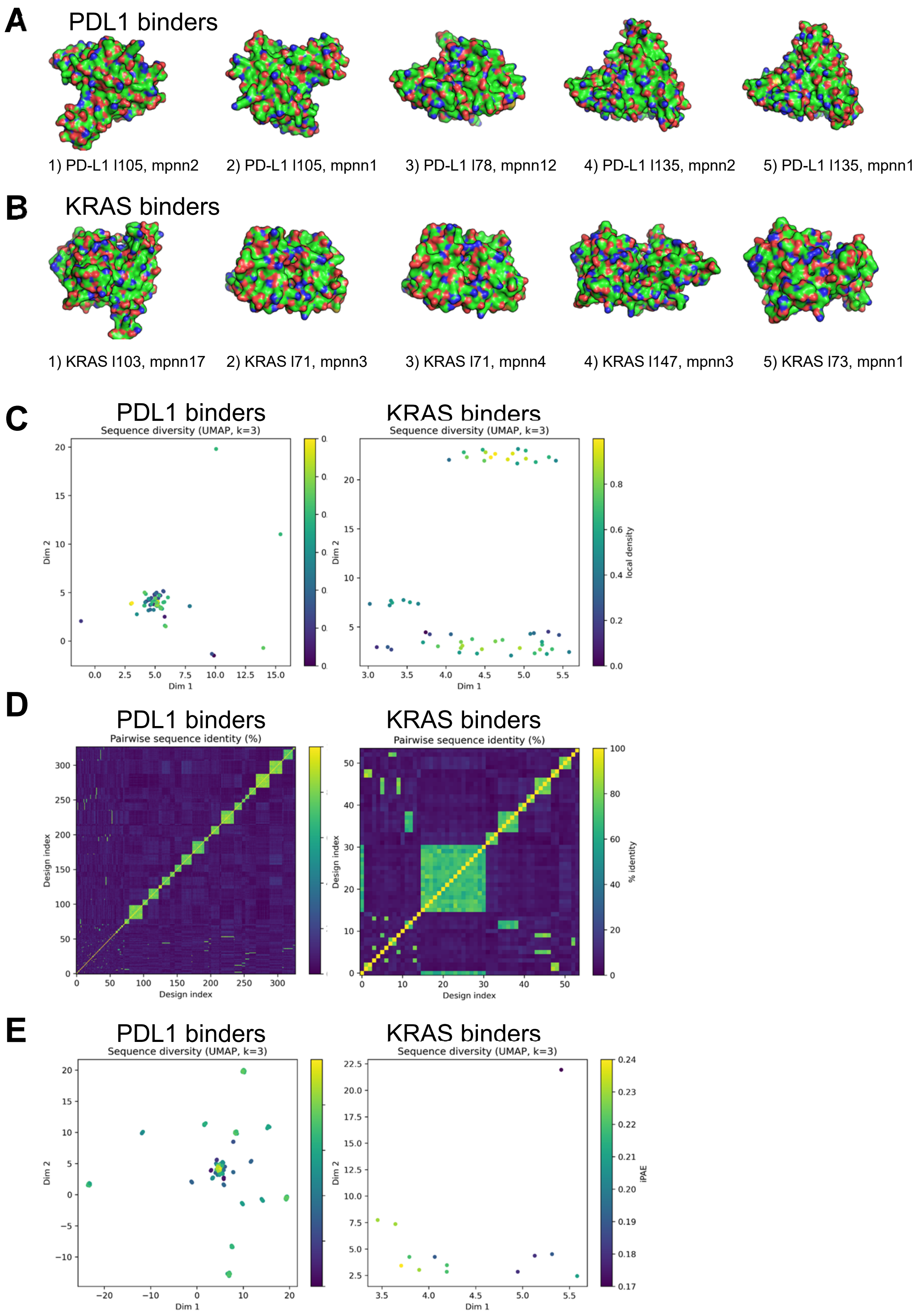

A
PDL1 binders
1) PD-L1 l105, mpnn2
2) PD-L1 l105, mpnn1
3) PD-L1 l78, mpnn12
4) PD-L1 l135, mpnn2
5) PD-L1 l135, mpnn1
B
KRAS binders
1) KRAS l103, mpnn17
2) KRAS l71, mpnn3
3) KRAS l71, mpnn4
4) KRAS l147, mpnn3
5) KRAS l73, mpnn1
C
PDL1 binders
Sequence diversity (UMAP, k=3)
KRAS binders
Sequence diversity (UMAP, k=3)
Dim 1
Dim 2
local density
D
PDL1 binders
Pairwise sequence identity (%)
KRAS binders
Pairwise sequence identity (%)
Design index
% identity
E
PDL1 binders
Sequence diversity (UMAP, k=3)
KRAS binders
Sequence diversity (UMAP, k=3)
Dim 1
Dim 2
iPAE


**Figure S3**

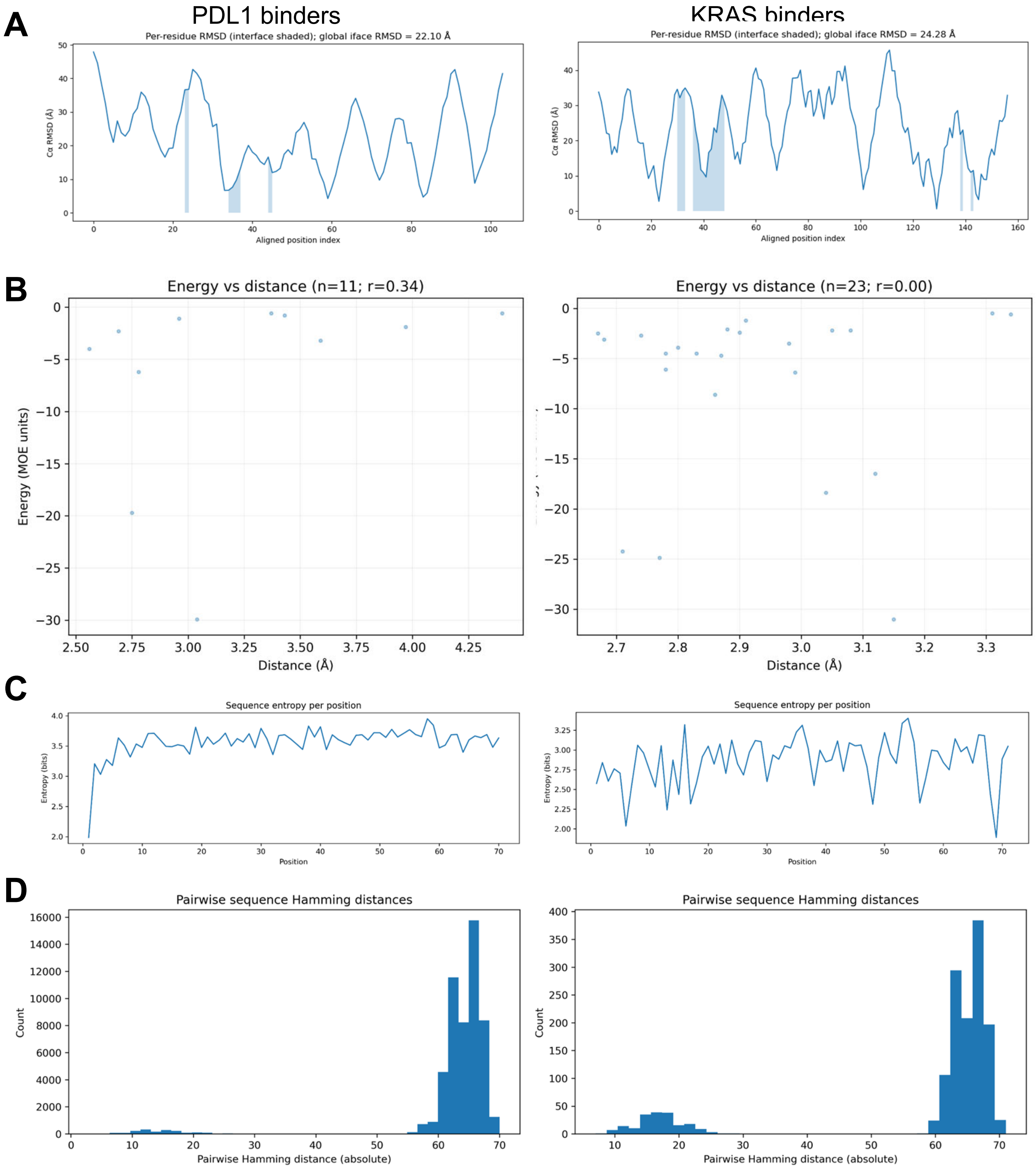
A
PDL1 binders
KRAS binders
Per-residue RMSD (interface shaded); global iface RMSD = 22.10 Å
Per-residue RMSD (interface shaded); global iface RMSD = 24.28 Å
Cα RMSD (Å)
Aligned position index
B
Energy vs distance (n=11; r=0.34)
Energy vs distance (n=23; r=0.00)
Energy (MOE units)
Distance (Å)
C
Sequence entropy per position
Entropy (bits)
Position
D
Pairwise sequence Hamming distances
Count
Pairwise Hamming distance (absolute)

Figure S4
